# The Distributed Engram

**DOI:** 10.1101/583195

**Authors:** Ran Feldesh

## Abstract

Neural connectionism is a common theoretical abstraction of biological neural networks (1–3) and a basis for common artificial neural networks (4). Yet, it is clear that connectionism abstracts out much of the biological phenomena significant and necessary for many cognitive-driven behaviors, in particular intra-neuronal and inter-neuronal biochemical processes (5–8). This paper presents a model which adds an abstraction of these processes to a standard connectionism-based model. Specifically, a sub-system determines the synaptic weights. The resulting network has plastic synapses during non-learning-related behavior, in sharp contrast with most common models in which synapses are fixed outside of a learning-phase. Some synapses introduce plasticity that is causally related with behavior, while in others the plasticity randomly fluctuates, in correspondence with recent data (9,10). In this model the memory engram is distributed over the biochemical system, in addition to the synapses. The model yields better performance in memory-related tasks compared to a standard recurrent neural network trained with backpropagation.

## Introduction

The biology underlying learning and memory is intricate. Computational models attempt to abstract out as many biological details as possible, while capturing functionality and maintaining a faithful representation of the neurobiology.

A common neuroscientific framework for learning and memory is neuronal connectionism. Most connectionist models assume that memory corresponds to a set of synaptic weights and learning is manifested by changes to these weights. The ‘synaptic plasticity and memory hypothesis’ states that: *“activity-dependent synaptic plasticity is induced at appropriate synapses during memory formation and is both necessary and sufficient for the information storage underlying the type of memory mediated by the brain area in which that plasticity is observed.”* (11–13).

In an integrate-and-fire connectionist model the parameters of the system are the synaptic weights, neuronal threshold and electrical activity. The dynamics is commonly divided into two distinct phases: learning, in which the weights are being changed, and learned behavior, in which the weights are constant (2,14,15). A similar view prevails in the standard formulation of artificial neural networks, where the synapses are plastic during learning, via a backpropagation process, and are fixed during ‘behavior’, such as prediction (4).

There is ample evidence supporting neuronal connectionism. One of the prominent findings is the discovery by Bliss and Lomo in the early 1970s that long-term potentiation (LTP) may serve as a substrate for long term memory (16). More recently, learning has been demonstrated to be correlated with a transient increase in the density of dendritic spines (17,18), and was shown to be necessary for memory formation (19). Spike-timing-dependent plasticity has been described as a learning mechanism (20).

However, additional findings correspond with a richer picture of the biological substrate of learning and memory; while the sufficiency of synaptic weights for memory might be true to a good approximation, in some cases, the neural repertoire is richer. First, synapses, in some cases, fluctuates without any apparent learning taking place, to a level similar to that observed during learning (9,21–24). This raises the question of locating the memory engram; if the synapses are dynamic and memory is preserved, then where is it stored?

Secondly, the dynamical processes related to memory dynamics are governed by biochemical processes, such as protein networks and chromatin modifications. Some of these processes take part as the neural and molecular basis of behavior, in addition to learning (25–29). Thirdly, neurons interact by exploiting dozens of neuromodulators, neuropeptides, diffusible molecules, hormones and RNA, whose effect is not restricted to learning (30–41).

Synaptic weights and electric currents, in these cases, are only part of a broader mechanism, and are not sufficient for memory induction, storage and expression. In these cases, a hypothetically complete knowledge of the synaptic weights is not sufficient to infer all memories within that system; a complete theoretical knowledge of the synaptic state, electrical activity and environmental input is not sufficient to predict behavior.

How can the additional observations of synaptic dynamics and the new molecular data be incorporated in a model that will also be simple enough to effectively capture very general aspects of learning and memory? This paper focuses on a single central question - where can the engram be located in a case of volatile synapses, given the role of the biochemical intraneuronal and interneuronal networks?

To answer that, a conceptual model is introduced, which adds a high-level representation of intraneuronal and interneuronal biochemical networks to a typical connectionist model. The intra-neuronal networks communicate through biochemical signals, such as neuromodulators and peptides, and together govern the weights of the synaptic network. The weight of each synapse in the output, or outer, network is a linear combination of the activity of the governing network. The governing network is represented by a neural network as such a formulation has been used to describe to a first approximation protein networks, gene networks and inter-neuronal biochemical networks (42–44).

While there are biological characteristics that this model lacks, such as absence of time integration and no segregation into separated internal and neuro-specific networks, it captures the essence of a system that continuously controls the synapses, hence allowing the engram to partly reside in that biochemical system. The model demonstrates that the memory engram can be distributed between synapses and intra- and inter-neuronal biochemical components. It predicts that in the case of volatile synapses, the engram can be stored in the biochemical networks. It does not point to specific pathways for this storage capacity, since it does not directly represent any specific pathway. The model only offers the high-level view that memory is distributed between synapses and their governing biochemical network. This view differs from other hypotheses that address the location of the engram given volatile synapses, which assume that memory resides in inhibitory synapses (45) or is represented as time-varying attractors in neural state-space (46). The suggestion presented in this paper does not negate these proposals, which may all co-exist.

To summarize, this model corresponds with, and highlights the following ideas:

1. Synaptic plasticity takes place in cases were no evident learning takes place.
2. The network dynamics during behavior includes in some cases biochemical changes in addition to electrical activity. Intraneuronal and interneuronal processes are crucial parts of the setting of behavior.
3. As a result of the above, induction, maintenance and expression of a memory depend not just on electrical activity and synaptic efficacy, but on biochemical interneuronal and intraneuronal processes.

In essence, it is suggested that synaptic weights are part of a larger mechanism that includes intraneuronal and interneuronal biochemical processes. Synaptic weights are not a sufficient component for memory in many biological cases. Rather, they can vary outside of a learning-related case. This is sharp contrast with many common models. In this case, the engram is distributed within the synapses and the biochemical network, as demonstrated in this model.

### The model

The standard description of an artificial neural network (ANN) consists of a set of interconnected processing elements, or neurons. It can be described as a directed graph in which each node is a function:

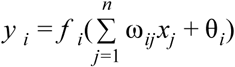

where *y*_*i*_ is the output of the node *i, x*_*j*_ is the *j*^*th*^ input to the node, and *w*_*ij*_ is the connection weight between node *j* to *i*. θ_*i*_ is the threshold (or bias) of the node. Usually, *f_i_* is a nonlinear function, such as a sigmoid or a linear-rectified function (47) ^2^ (figure 1).

**Figure 1.**
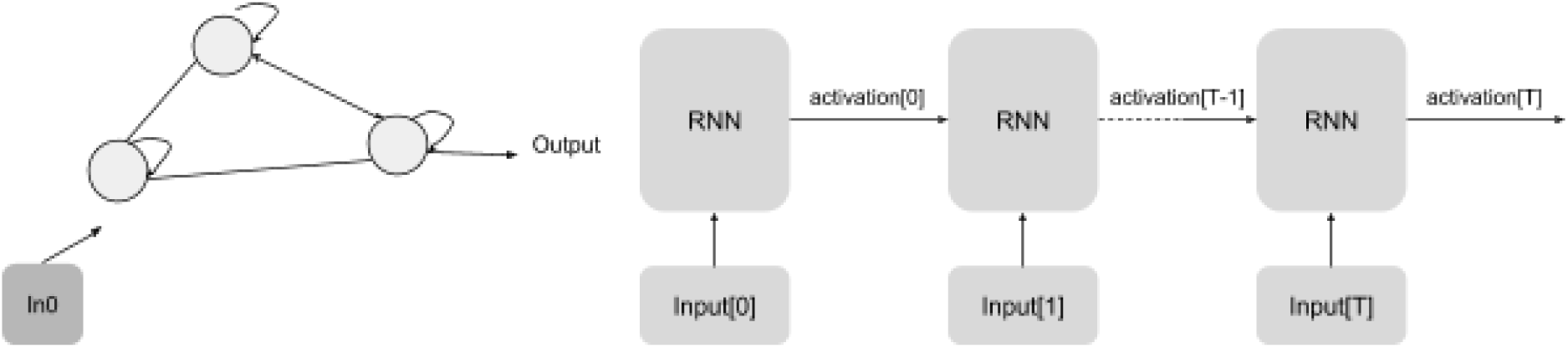
A standard RNN. Left: a fully connected network of three neurons. Right: a RNN over *T* time steps (three are shown), with the input and activations. The weights and biases are optimized during training, and stay fixed afterwards.

The plastic Recurrent Neural Network (RNN) model, includes an inner, neural network, which continuously sets the weights of the outer network during prediction, or behavior. The output of the system is the value of one of the neurons of the outer neurons, while the inner network’s output is latent (hence the naming, ‘outer’ and ‘inner’). This is represented by the following equations:

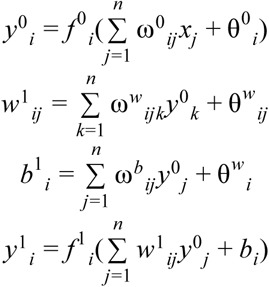

The 0 and 1 superscripts stands for the inner and outer networks respectively, so that *y*^0^ and *f* ^0^ are the activation value and function of the inner network, and *y*^1^ and *f* ^1^ are corresponds to the outer one. Importantly, only ω^0^, θ^0^, ω^*w*^, θ^*w*^, ω^*b*^, θ^*b*^ are trainable parameters (i.e., they are the only parameters changed during the backpropagation process). The weights and threshold of the outer network, *w*^1^ and *b*^1^ are dynamically set by the activation of the inner network. The external input flows to both the inner and the outer networks (figure 2).

**Figure 2.**
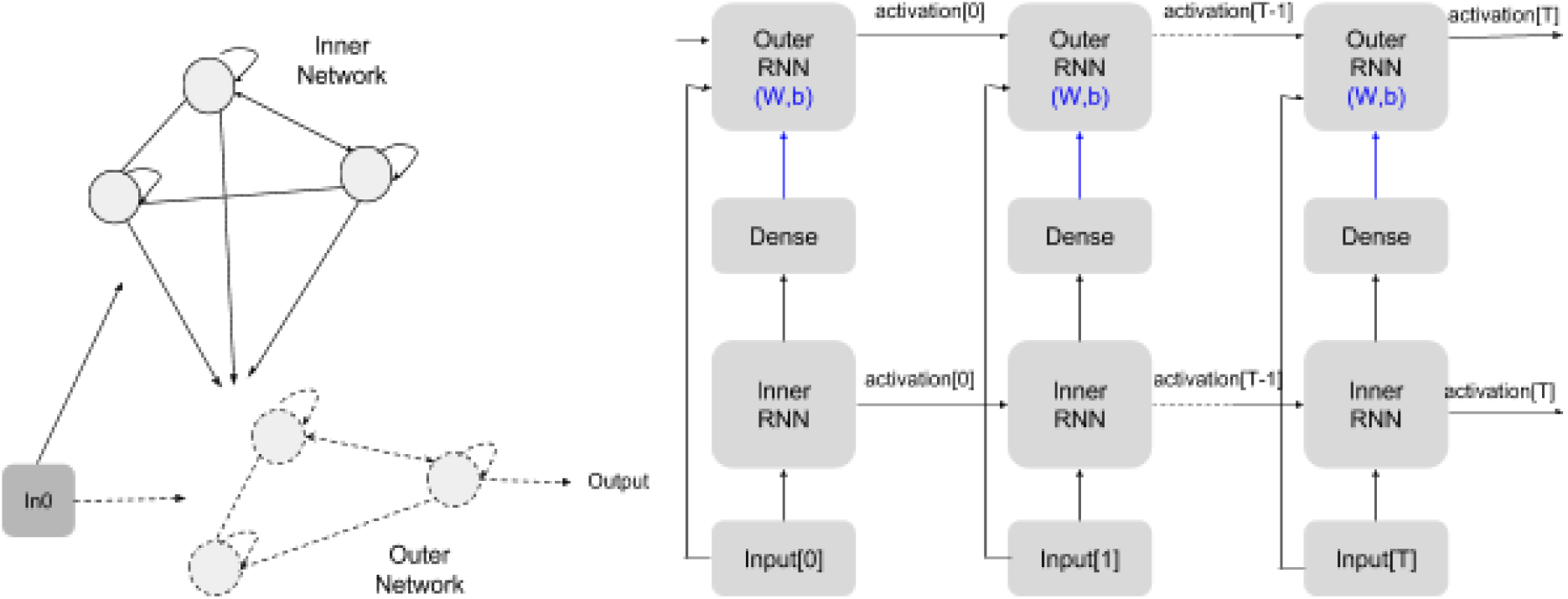
A plastic RNN. Left: a neural network with three neurons, governed by a second network with three interacting factors. The governing network receives external input, and its output sets the weights of the main network. Right: the plasticRNN over *T* timesteps (three are shown). The inner network’s parameters (left: solid black lines; right: blue) are being optimized during training and are kept fixed during prediction (behavior). The outer networks’ weights and biases, are being determined by the inner network activity.

### Binary classification task

A boolean function, such as an AND, OR, XOR, NOR, defines a 2d to 1d mapping, and can be generalized to a mapping over an *n*-dimensional space. For instance, a two-arguments AND function is a boolean rule with four optional inputs: [0, 0], [0, 1], [1, 0], [1, 1], which maps to 0, 0, 0, 1 accordingly. The complexity of the problem partly depends on the dimensionality of the input vector. Here, *n* = 2.

The binary classification task is a time-sequence classification task. Each trail is a sequence of *T* timesteps. Each timestep is associated with a boolean rule. The rule is fixed for a given number of time-steps, and then changes to another, and so forth until the end of the trail.

The input at each time-step is a vector of *n* + 1 binary elements, where *n* is the input to the boolean function (*n* = 2). The additional input at time *t* is the result of the boolean rule at *t* – 1 with the input of *t* - 1. Therefore, at each timestep the network receives a feedback in the form of delayed ground truth (DGT) of the previous time-step. At each time-step, the network output is a scalar within the range [0, 1]. The goal of the network is to respond, at every timestep, in accordance to the binary rule at that timestep.

For example, figure 3 describes a single trail, with 20 timesteps, while each timestep has three inputs. The first 9 timesteps correspond to an AND binary rule, a correct answer by the network would be 1 for *t* = 3, 5, 6 and 0 the otherwise. Likewise, timesteps 9-19 satisfy the OR binary rule. A correct answer would be 0 for *t* = 11, 13, 18 and 1 otherwise. At every timestep during both training and testing the network gets a delayed ground truth with the correct answer of the previous timestep.

**Figure 3.**
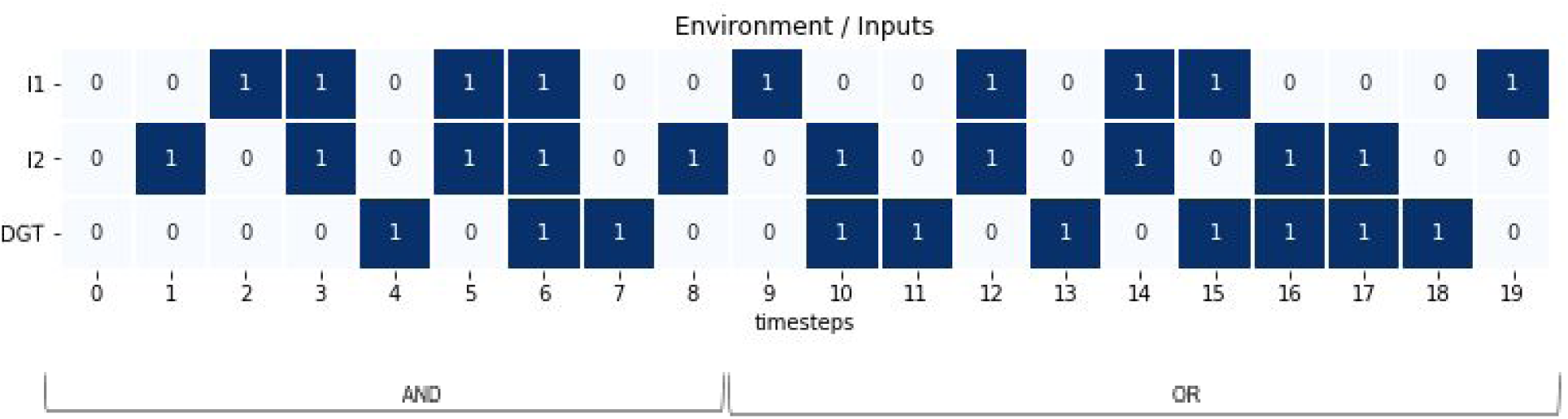
An example of a trail. Each trail is composed of a sequence of 3D inputs and, in this case, 20 time-steps. Each time-step is associated with a Boolean rule, shown at the below the x-axis; an AND rule takes place between time-steps 0 to 8, while an OR rule is between time-step 9 to 19. At time-step *t* the network’s response is correct if it is identical to the outcome of the boolean rule for the inputs, *I*1 and *I*2, at that timestep. *DGT* is the Delayed Ground Truth: at timestep *t* it indicates the correct response for the previous timestep, *t* − 1. For any sequence, the DGT at *t* = 0 is arbitrarily set to zero. The DGT for the last timestep is not given to the network.

A two-stage solution might be to first identify the binary rule, and then respond accordingly. Clearly, the actual prediction by the network might not follow this procedure, however, a memory of the current rule has to be stored by the network, as it is not possible to know the rule based on any single time-step.

## Results

In plasticRNN, the activation, weights and biases of the outer network all change dynamically during prediction. This is demonstrated in figure 4. The accuracy of plasticRNN was calculated following training for a range of inner and outer network sizes. Trails containing 12 binary rules were chosen for training and testing, and the duration of each rule was fixed to 30 time steps. Accuracies larger than 90% were seen for most networks where the number of trainable parameters was larger than 200 (figure 5).

**Figure 4.**
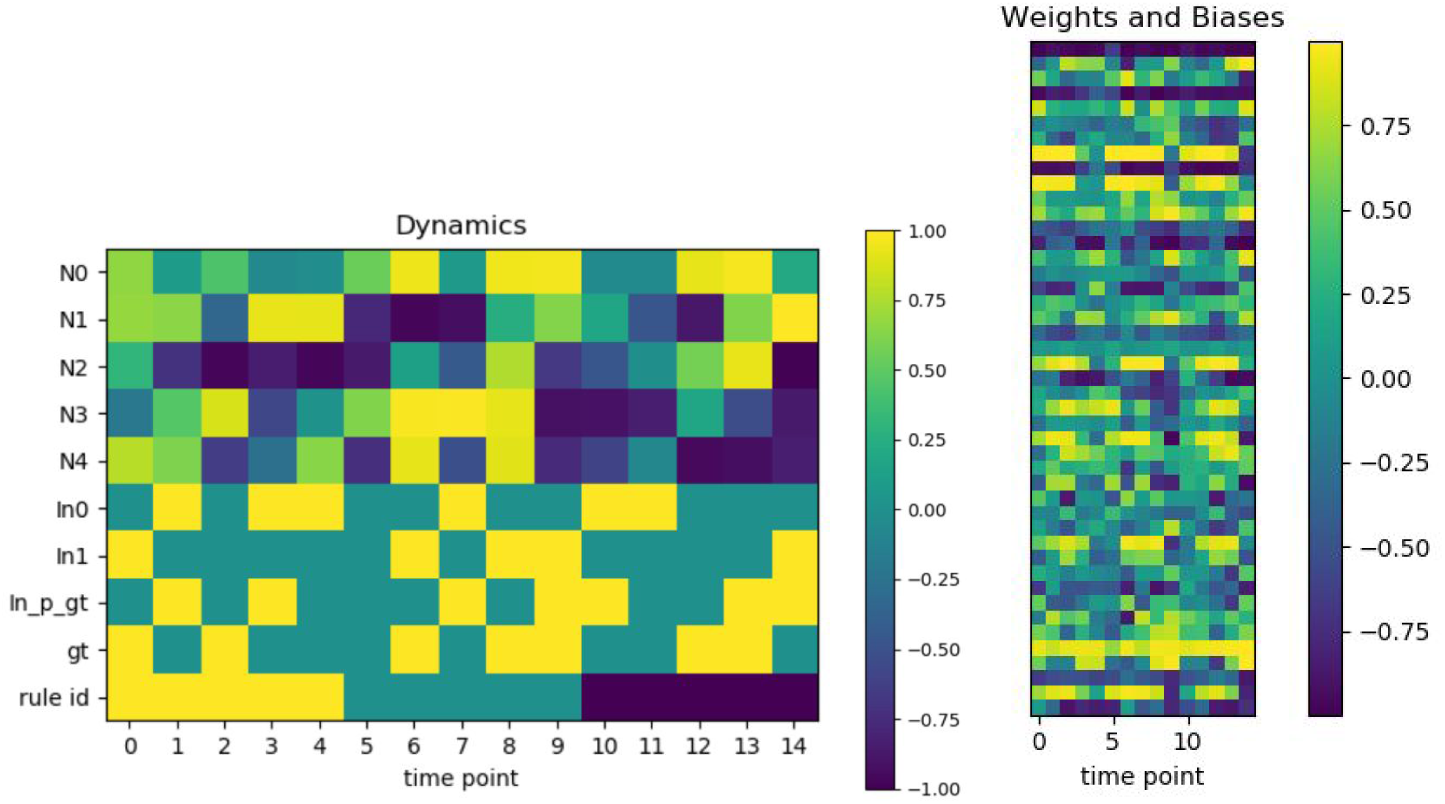
left: Example of the dynamics of the activations for a plasticRNN, with five neurons (N0-N4) in the outer network. The three inputs (*In*0, *In*1 and *In*_*p*_ *gt,* the delayed ground truth) appear next. *gt* is the ground truth at that time-step. Three rules, each lasting for 5 time-steps compose this trail. The *gt* and rule id are not presented to the network. Right: The dynamics of the weights and biases of the outer network during the trail.

**Figure 5.**
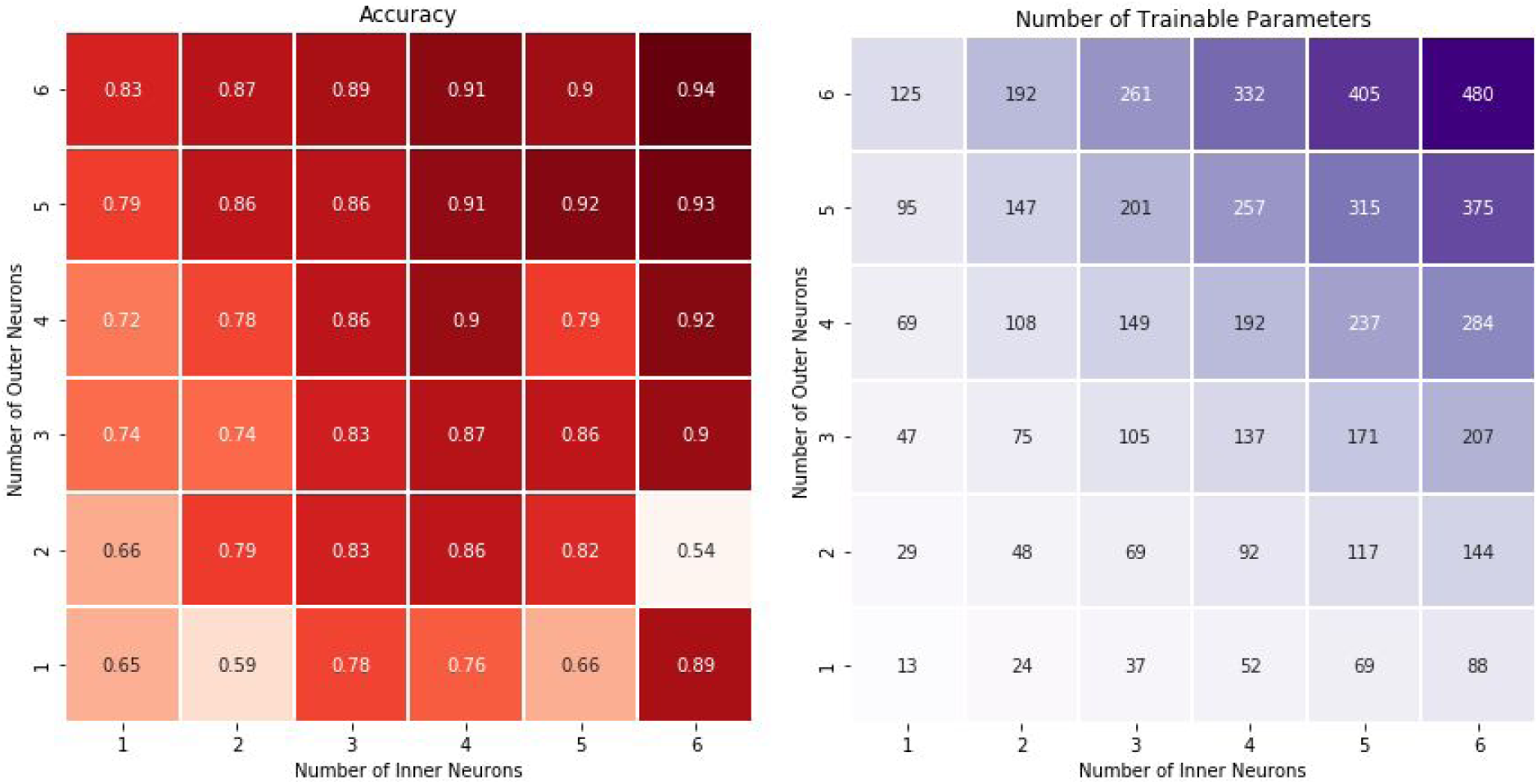
Left: Average accuracy over 10 sets of training and testing. The network size was varied within the range [1-6] for the inner and outer network size. Each trail is composed of 12 rules, each for 30 timesteps. Right: number of trainable parameters of a plasticRNN as function of the number of inner and outer neurons.

Next, the performance of plasticRNN was compared to that of a standard RNN (BasicRNN). The plasticRNN accuracy was above 94%, on average, for sequences with two to ten rules. Comparably, while the standard RNN performance started at 93% for sequences of two and three rules, it degraded to 72-78% for sequences with five or more rules (figure 6). PlasticRNN was compared to state of the art networks, LSTM and GRU, as well as to multi-layered RNN. PlasticRNN performs similarly to these networks (figure 7). The number of trainable parameters was comparable between all these networks.

**Figure 6.**
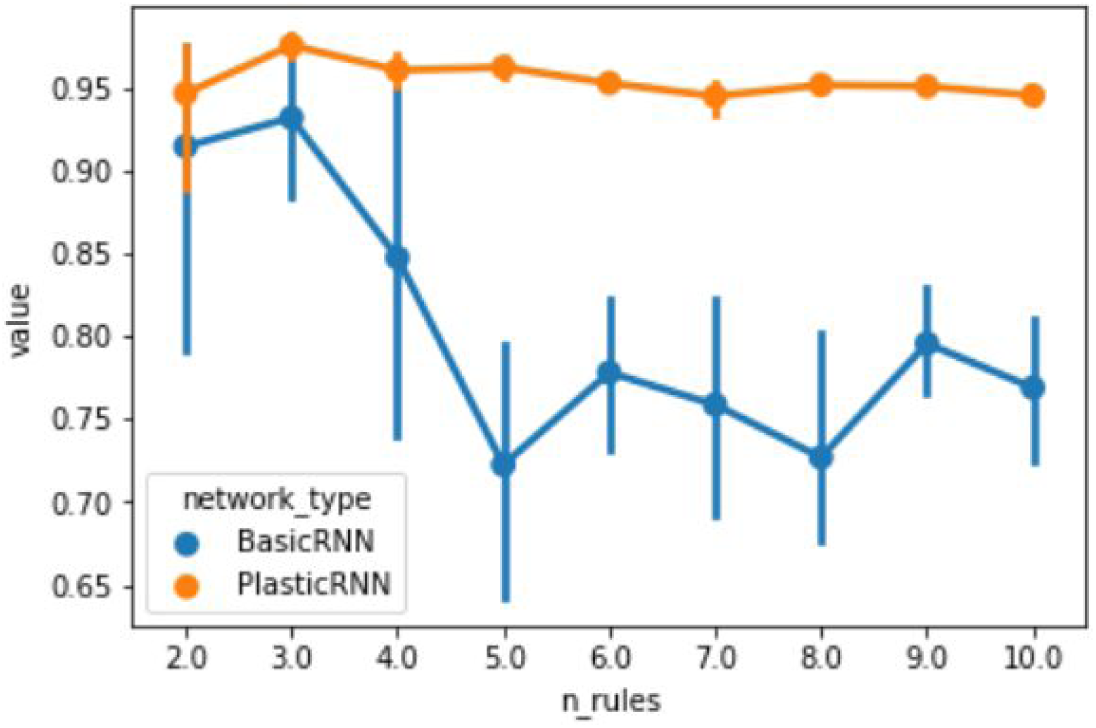
A standard RNN (BasicRNN) and a PlasticRNN have been trained and tested with sequences of varying number of rules (n_rules). The average accuracy and standard deviation over ten independent tests for each value of n_rules are presented.

**Figure 7.**
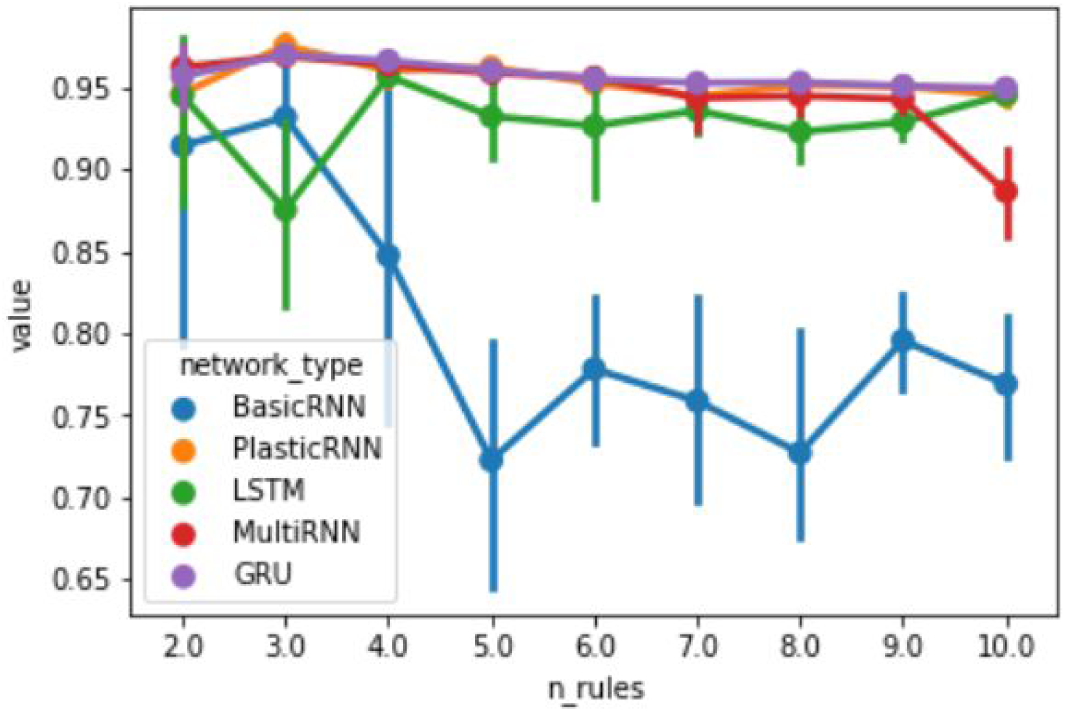
Comparison of plasticRNN with a standard RNN (BasicRNN), LSTM, Multi layered RNN and GRU, for sequences with varying number of rules (n_rules).

Importantly, the weights of the outer network have been compared across four rules, for a small plasticRNN network, with three neurons in its outer network. The weights of the outer network had distinct values for distinct rules: 14 out of the 18 weights, 78%, had unique values per rule.

**Figure 8.**
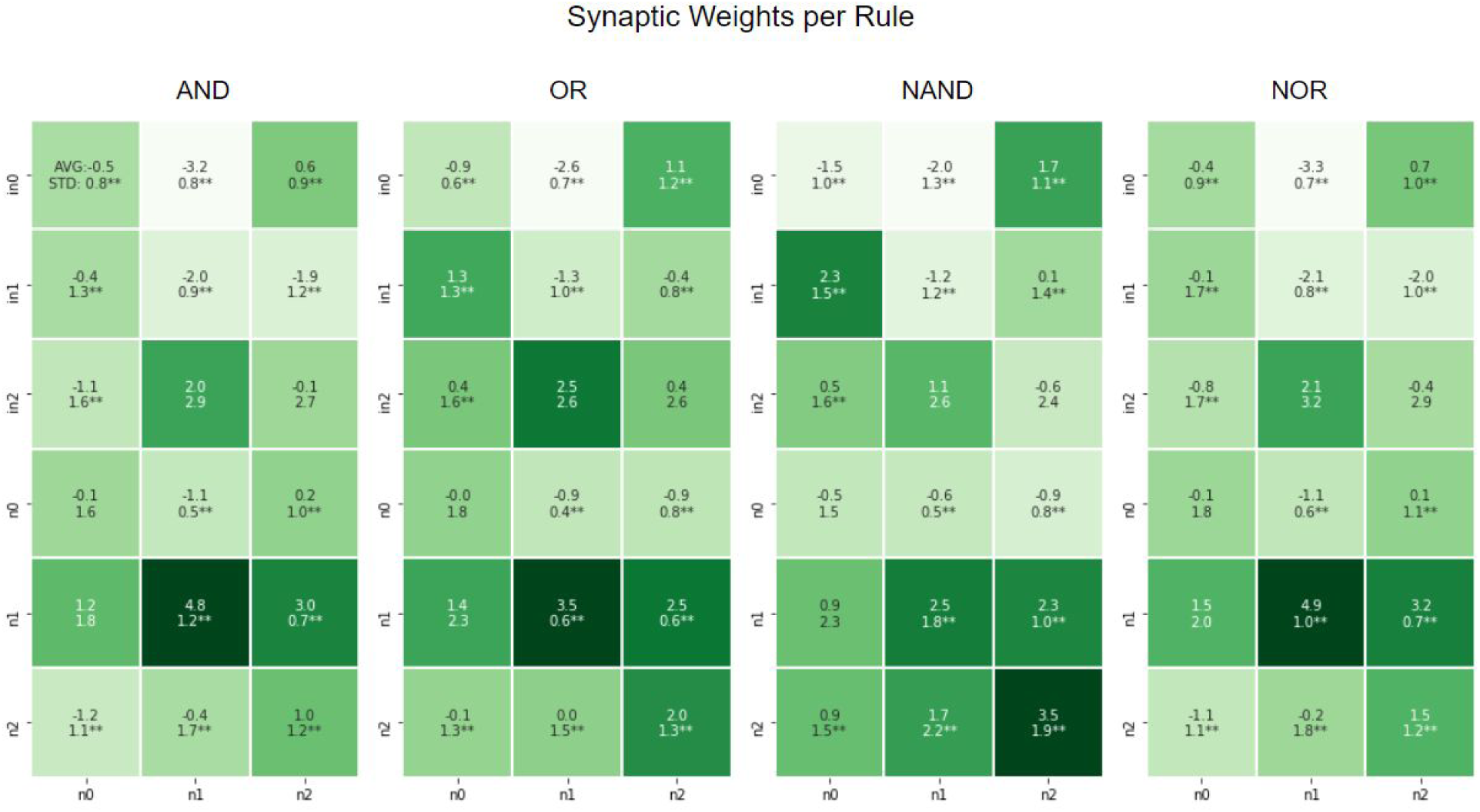
Synaptic weights per Boolean rule during prediction, at test time, post training, for a network with 3 external neurons. The network was trained on 12 rules. Average and standard deviation of each plastic weight within each rule are presented. A double asterisk represents a Bonferroni-corrected p value < 0.01.

## Discussion

This paper presents a neural network model which addresses the question of the location of the engram in a case of volatile synapses (9). The model adds to a standard connectionist models an abstract representation of the intraneuronal and neuromodulatory interactions which together control the synaptic weights. A simple model, in which one neural network sets the weights of another one during prediction, performs better than a standard recurrent neural network, when both are trained with backpropagation, in a memory-related task. Moreover, the synaptic weights fluctuate during behavior and not just during learning. Importantly, many of the fluctuations are correlated with behavior, and are necessary for successful performance. Therefore, in this model the memory engram does not reside just in the synaptic weights, but is distributed *also* within the system that governs them. We argue that a similar case, based on intracellular memory and neuromodulatory inter-neuronal signalling, occurs in biological systems.

The model is conceptual and simplistic. This is its power and its weakness. It captures two generic biological concepts, which are well-known but are not commonly modelled: *(a)* neural plasticity is not limited to the learning phase, and *(b)* that synaptic efficacies depend on intra- and inter-neuronal processes in a non-trivial way. In nature, learning is manifested by multiple biochemical processes, which a single algebraic equation (e.g. Hebb rule or spike-time dependent plasticity) does not fully capture. The weakness point of this model is in the details: the elements of the governing-network do not directly correspond to identified intra- and inter-neuronal biological entities.

The simplicity of the model is appealing and valuable in conveying the idea of an engram that is diffused in both the synapses *and the biochemical network that governs them*. This is in contrast to other hypotheses addressing the engram location given volatile synapses, such as having the memory in inhibitory synapses (45) or as time-varying attractors in neural state-space (46), although all these mechanisms can co-exist. Future directions include introducing segregation and time delays into the governing network, and extending the test to additional tasks.

## Acknowledgment

*I thank Eva Jablonka for her useful comments. Any error is mine.*

## Footnotes

2 While this is by far the most common description of an ANN, a more general case is sometimes considered, see for example (14), p.46 and p.51.

